# The conserved global regulator H-NS has a strain-specific impact on biofilm formation in *Vibrio fischeri* symbionts

**DOI:** 10.1101/2024.12.19.629378

**Authors:** Dani Zarate, Ruth Y. Isenberg, Morgan Pavelsky, Lauren Speare, Aundre Jackson, Mark J. Mandel, Alecia N. Septer

## Abstract

Strain-level variation among host-associated bacteria often determines host range and the extent to which colonization is beneficial, benign, or pathogenic. *Vibrio fischeri* is a beneficial symbiont of the light organs of fish and squid with known strain-specific differences that impact host specificity, colonization efficiency, and interbacterial competition. Here, we describe how the conserved global regulator, H-NS, has a strain-specific impact on a critical colonization behavior: biofilm formation. We isolated a mutant of the fish symbiont *V. fischeri* MJ11 with a transposon insertion in the *hns* gene. This mutant formed sticky, moderately wrinkled colonies on LBS plates, a condition not known to induce biofilm in this species. A reconstructed *hns* mutant displayed the same wrinkled colony, which became smooth when *hns* was complemented *in trans*, indicating the *hns* disruption is causal for biofilm formation in MJ11. Transcriptomes revealed differential expression for the *syp* biofilm locus in the *hns* mutant, relative to the parent, suggesting biofilm may in part involve SYP polysaccharide. However, enhanced biofilm in the MJ11 *hns* mutant was not sufficient to allow colonization of a non-native squid host. Finally, moving the *hns* mutation into other *V. fischeri* strains, including the squid symbionts ES114 and ES401, and seawater isolate PP3, revealed strain-specific biofilm phenotypes: ES114 and ES401 *hns* mutants displayed minimal biofilm phenotypes while PP3 *hns* mutant colonies were more wrinkled than the MJ11 *hns* mutant. These findings together define H-NS as a novel regulator of *V. fischeri* symbiotic biofilm and demonstrate key strain specificity in that role.

**Importance:** This work, which shows how H-NS has strain-specific impacts on biofilm in *Vibrio fischeri*, underscores the importance of studying multiple strains, even when examining highly conserved genes and functions. Our observation that knocking out a conserved regulator can result in a wide range of biofilm phenotypes, depending on the isolate, serves as a powerful reminder that strain-level variation is common and worthy of exploration. Indeed, uncovering the mechanisms of strain-specific phenotypic differences is essential to understand drivers of niche differentiation and bacterial evolution. Thus, it is important to carefully match the number and type of strains used in a study with the research question to accurately interpret and extrapolate the results beyond a single genotype. The additional work required for multi-strain studies is often worth the investment of time and resources, as it provides a broader view of the complexity of within-species diversity in microbial systems.

## Results

Strain-level variation within bacteria has been observed across diverse species and can influence a wide range of ecological functions including host range and disease (1-4). *Vibrio fischeri* is a bioluminescent, beneficial symbiont that colonizes the light organs of fish and squid (5). The association between *V. fischeri* and *Euprymna scolopes* squid has served as a valuable model system for studying the genetic determinants and molecular mechanisms underlying beneficial associations between bacteria and animals (6). Previous studies have described strain-level differences among *V. fischeri* isolates in which conserved functions or genes are differentially regulated (7). A few notable strain-specific differences include bioluminescence output and regulation (8), biofilm formation (9, 10), and interbacterial killing via the type VI secretion system (11, 12).

Here, we explore the extent to which a conserved regulator (H-NS) has strain-specific effects on a conserved behavior in *V. fischeri*: biofilm formation. H-NS is a global regulator of gene expression during environmental transitions (13) and has been previously connected to biofilm in other species including *Aggregatibacter actinomycetemcomitans* (14), *Vibrio cholerae* (15), and *Klebsiella pneumoniae* (16). We begin by applying a combination of genetics, transcriptomics, microscopy, and host colonization assays to a model *V. fischeri* strain, MJ11, and then determine the extent to which H-NS regulates biofilm in *V. fischeri* strains from more diverse isolation sources.

### The MJ11 *hns*::tn5 mutant has wrinkled colony morphology

Previously, we conducted a random transposon mutant screen in the fish symbiont *V. fischeri* MJ11 (17) and noticed one mutant produced sticky, wrinkled colonies on plates (Fig 1A).This mutant had a transposon inserted into VFMJ11_1751 (LAS35E11), which encodes for the global regulator H-NS. The protein is an ortholog of characterized *V. cholerae* H-NS VC_1130, with 61 % identity and 73 % similarity across the entire protein. To verify the biofilm phenotype was due to the *hns* disruption, and not a secondary mutation, we used natural transformation to move the mutation in LAS35E11 into a fresh MJ11 background, resulting in strain DZ101, which also produced wrinkled colonies. When *hns* was complemented *in trans* by introducing plasmid pNL6 (18) into DZ101, the complemented strain produced smooth colonies (Fig 1A).

**Figure 1.**
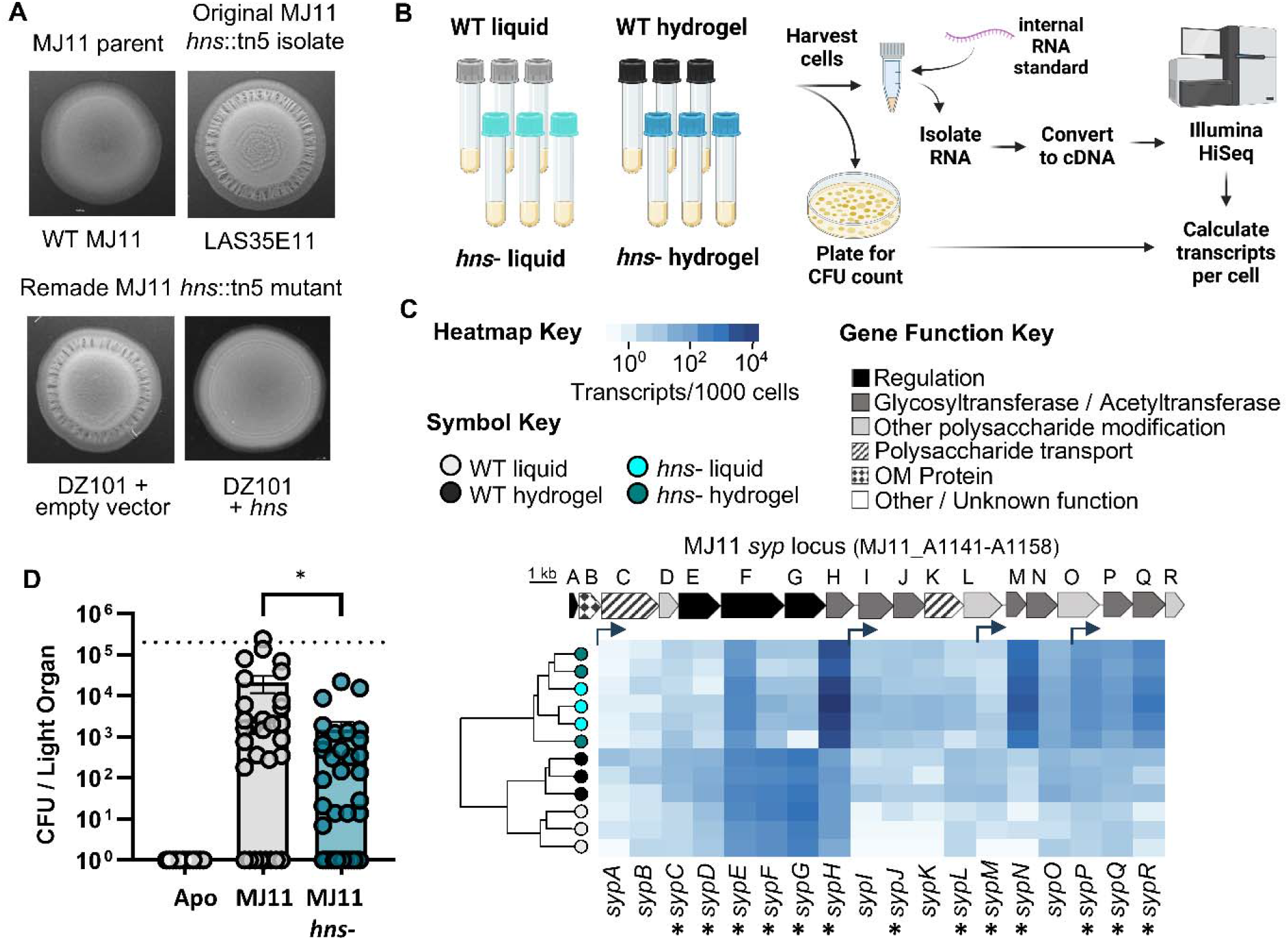
The *hns* mutation derepresses biofilm in MJ11 but does not show an increased colonization ability. (A) LAS35E11 or remade MJ11 *hns*::tn5 mutant (DZ101) with empty vector (pVSV105) or *hns* complementation vector (pNL6). Representative images are after 48 hr incubation on LBS plate at 24C. All images were captured using a Leica M165 FC microscope with Flexcam C3 camera. Images were converted to grayscale and brightness and contrast were adjusted uniformly. (B) Methods flowchart for obtaining quantitative transcriptomes. Made with Biorender.com. (C) Heatmap of hierarchical clustering results for the *syp* (VFMJ11_A1141-A1158) and *bcs* (VFMJ11_A1000-A1007) gene clusters indicating transcripts per 1000 cells for MJ11 wild-type (WT) grown in liquid (gray) or hydrogel (black) and *hns-* mutant grown in liquid (cyan) in liquid or hydrogel (dark cyan). Bent arrows indicate predicted promoter locations based on work in ES114. Each row represents a sample and each column represents a gene; gene ID is shown at the bottom of the lower heatmap in each panel. Square color in the heatmap indicates the absolute abundance of each transcript per cell. Asterisks indicate statistically significant differences comparing WT and *hns-* in hydrogel (I-test, p<0.05). (D) CFU per light organ for each animal. Animals were exposed to indicated inoculum for 3 hr. CFUs obtained at 48 hr post inoculation. Dashed line indicates average level of colonization for native *V. fischeri* symbionts. Data are combined results from four separate experiments with a total of animals for each treatment being: 12 (Apo), 30 (MJ11), and 36 (MJ11 *hns* mutant LAS35E11).

### Transcriptomic analysis reveals changes in *syp* biofilm gene expression in the *hns* mutant

To better understand how gene expression changes might impact biofilm formation in the MJ11 *hns* mutant, we performed a quantitative transcriptome analysis for wild-type and *hns* mutant cultures grown in liquid LBS or hydrogel (LBS supplemented with 5% w/vol polyvinylpyrrolidone, PVP) (Fig 1B). A hierarchical cluster analysis showed that the *syp* locus, which encodes factors that produce the SYP polysaccharide required for biofilm and aggregate formation during colonization of *Euprymna scolopes* squid (19), displayed significant differences in gene expression that grouped by genotype (Fig 1C). Specifically, the *hns* mutant cultures in both conditions showed increased expression of genes predicted to be involved in symbiotic polysaccharide synthesis or modification, including up to 60-fold increases in transcript abundance for *sypH, sypN*, and *sypR* (Fig 1C, Table S1). We also examined expression of *bcs* genes that are responsible for cellulose production (20), and although we did not see significant differential expression across treatments, *bcs* genes were expressed in all strains and conditions and therefore the role of cellulose cannot be ruled out as part of the biofilm mechanism in the *hns* mutant.

### Biofilm production in the MJ11 *hns* mutant is not sufficient to permit host range expansion

Given that MJ11, a fish isolate, does not efficiently colonize the *E. scolopes* squid host unless biofilm is induced (9, 21), we asked whether the MJ11 *hns* mutant might exhibit improved colonization. To answer this question, we exposed juvenile *E. scolopes* squid to wild-type MJ11 or the MJ11 *hns* mutant and compared colonization levels as a measure of CFUs per animal at 48 hours post inoculation. Despite the biofilm phenotype in culture, the MJ11 *hns* mutant was even less effective at colonizing the squid than the wild-type parent (Fig 1D), indicating that i) the biofilm production in the *hns* mutant is not sufficient to initiate symbiosis and/or ii) other symbiosis factors are negatively affected by the *hns* mutation. Indeed, Lyell *et al*. observed a colonization defect for an *hns* mutant of strain ES114 (22), a natural symbiont of the squid.

### H-NS has a strain-specific impact on biofilm formation

Given that *hns* and biofilm are both conserved across *V. fischeri* strains, we asked whether the *hns* mutation might similarly induce biofilm formation in diverse isolates. To answer this question, we used natural transformation to move the *hns*::tn5 mutation from LAS35E11 into two *E. scolopes* light organ isolates (ES114 and ES401) as well as the seawater isolate PP3, resulting in strains MP110, ANS3001, and MP111, respectively. We assessed the *hns* mutant and parent strains for biofilm in both surface and liquid-grown conditions using methods described in (23). Interestingly, both ES114 and ES401 *hns* mutants displayed relatively smooth colonies when grown on surfaces, while the PP3 *hns* mutant was highly wrinkled (Fig 2A), even more so than the MJ11 *hns* mutant. When we assessed biofilm growth on the surface of standing liquid cultures, all three mutants appear to form a film of growth that could be observed when disrupted with a pipette tip (Fig 2B). Together, these results indicate that although H-NS appears to repress biofilm at least to some degree in the strains tested here, the strength of the phenotype varied widely across strains. It is worth noting that, while the H-NS homolog tested here is conserved (MJ11 and ES114 H-NS sequences share 100% identity), additional histone-like proteins are present in at least some of our tested strains (ex. MJ11_B0192), which could impact gene expression as in other species (24).

**Figure 2.**
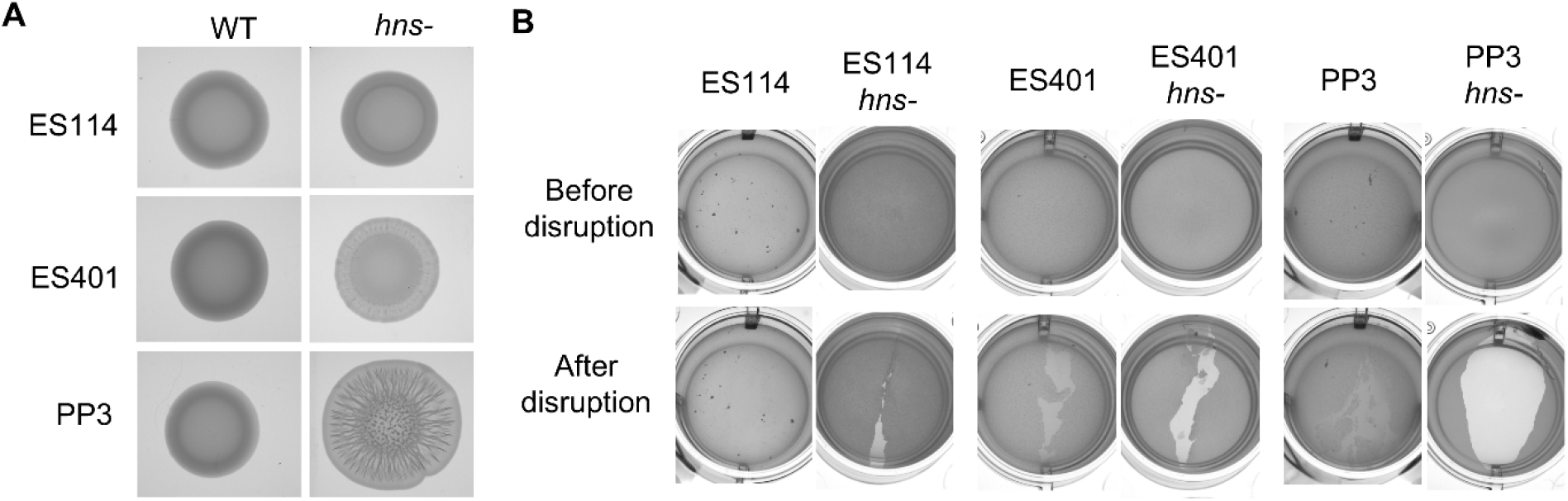
The effect of the hns mutation on biofilm phenotypes is strain specific. Representative of image wild-type (WT) ES114 and PP3 and their *hns* mutants after 48 hr incubation on LBS plates (A) or in standing culture (B) at 24° C. All images were captured using a Leica M165 FC microscope with Flexcam C3 camera

## Methods

See supplementary documents for details on methodology, strains and plasmids, quantitative transcriptomes, and imaging.

## Data Availability

Transcriptome data are available in supplemental files and via GenBank under BioProject ID PRJNA1013100.

## Acknowledgements

Work in the lab of Alecia Septer was supported by NIH NIGMS grant R35 GM137886 and Gordon and Betty Moore Foundation grant GBMF9328. LS was supported by a UNC dissertation completion fellowship. MJM was supported by NIGMS grant R35 GM148385. RYI was supported by NIGMS training grant T32 GM007215.

