## Supplemental Methods for "The conserved global regulator H-NS has a strain-specific impact on biofilm formation in *Vibrio fischeri* symbionts"

**Supplemental methods for Zarate et al.:**

**Bacterial strains, plasmids, and media.** *V. fischeri* strains were grown at 24-5°C in Luria-Bertani salt (LBS) medium (per liter: 25 g Difco LB broth [BD], 10 g NaCl, 50 mL 1 M Tris buffer [pH 7.5]). *E. coli* strains used for cloning and were grown at 37°C in Luria-Bertani (LB) medium (per liter: 25 g Difco LB broth [BD]). When needed, antibiotics were added to the media at the following concentrations: kanamycin, 100 μg/mL for *V. fischeri* and 50 μg/mL for *E. coli*; chloramphenicol, 1 or 5 μg/mL for *V. fischeri* and 25 μg/mL for *E. coli*; erythromycin, 5 μg/mL for *V. fischeri* and 150 μg/mL for *E. coli* grown in BHI medium. Growth media was solidified using 1.5% agar when needed.

**Isolation of LAS35E11.** The MJ11 *hns*::tn5 mutant (LAS35E11) was a false positive isolated during a transposon mutant screen to identify *V. fischeri* MJ11 mutants that lost the ability to kill target strains with their T6SS (Speare et al., 2024). The tn5 transposon on plasmid pEVS170 was introduced into MJ11 using triparental matings and ErmR transposon mutants were selected on LBS Erm plates as in (Lyell et al., 2008). Colonies were picked and grown in liquid LBS before screening for loss target killing. Transposon mapping was achieved using inverse PCR and Sanger sequencing and blasting results against the MJ11 genome (Mandel et al., 2009). The tn5 transposon inserted at bp 331 in the *hns* gene (VFMJ11_1751).

**Transcriptomes.** Quantitative transcriptomes were obtained for MJ11 WT and the LAS35E11 *hns*::tn5 mutant grown in liquid LBS (liq) or hydrogel, LBS with 5% w/v polyvinylpyrrolidone (PVP) as described in Speare et al., 2024. Transcriptome data are available in Table S1 and via GenBank under BioProject ID PRJNA1013100.

**Natural transformation.** To move the *hns*::tn5 mutation from MJ11 strain LAS35E11 to a fresh MJ11 background, or other *V. fischeri* isolates, genomic DNA was isolated from LAS35E11 using a Zymo Quick-DNA Fungal/Bacterial Miniprep Kit. Recipient strains MJ11, PP3, ES401, and ES114 were transformed with pLostfox-Kn and cells were prepared for natural transformation as in (Brooks et al., 2014). Briefly, a single colony of recipient strain was used to start a 3 mL LBS culture supplemented with Kanamycin. Cultures were grown overnight at 24°C with shaking. In the morning, 1 mL culture was centrifuged at 15,000 rpm for one minute, and resuspended in 1 mL Tris Minimal Medium (ref) supplemented with NAG and Kan. The cell suspension was then diluted 1:100 in 3 mL Tris Minimal Medium supplemented with NAG and Kan and grown to an OD600 of 0.2 - 0.5. Next, 500 ul of the prepared recipient culture was mixed with ~2.4 ug of donor DNA and allowed to incubate at room temperature for 30 min.  The mixture was moved to a fresh glass test tube, 1 mL LBS was added, and the cells recovered shaking overnight at 24°C. In the morning, the cells were plated onto LBS Erm plates to select for the mutation of interest (*hns*::tn5). Individual colonies were picked, restreaked and verified to have acquired the mutation (Erm-resistant) and lost the plasmid (Kan-sensitive).Strains and plasmids used in this study.

| **Strain** | **Relevant characteristics** | **Reference** |
| --- | --- | --- |
| ES114 | *V. fischeri;* isolated from *Euprymna scolopes* light organ | Boettcher and Ruby 1990 |
| ES401 | *V. fischeri;* isolated from *Euprymna scolopes* light organ | Fidiopastis et al, 2002 |
| PP3 | *V. fischeri;* Planktonic isolate, Kaneohe Bay, HI | Lee and Ruby, 1992 |
| MJ11 | *V. fischeri;* isolated from *Moncentris japonica* light organ | Ruby and Nealson 1976 |
| LAS35E11 | MJ11 *hns::tn5* mutant (Erm^R^) | This study |
| DZ101 | MJ11 *hns::tn5* remade from LAS35E11 gDNA (Erm^R^) | This study |
| DZ101 pVSV105 | MJ11 *hns::tn5* with empty vector (Erm^R^, Cm^R^) | This study |
| DZ101 pNL6 | MJ11 *hns::tn5* with ES114 *hns* complement plasmid (Erm^R^, Cm^R^) | This study |
| ANS3001 | ES114 *hns::tn5* made from LAS35E11 gDNA (Erm^R^) | This study |
| MP110 | ES401 *hns::tn5* made from LAS35E11 gDNA (Erm^R^) | This study |
| MP111 | PP3 *hns::tn5* made from LAS35E11 gDNA (Erm^R^) | This study |
| *E. coli* CC118λpir | *E. coli; Δ(ara-leu) araD Δlac74 galE galK phoA20 thi-1 rpsE rpsB argE*(Am) *recA λpir* | Herrero *et al.,* 1990 |
| *E. coli* DH5⍺λpir | *λpir derivative of E. coli; F’/endA1 hsdR17 glnV44 thi-1 recA1 gyrA relA1* Δ*(lacIZYAargF)*  *U169deoR(f80dlacI* Δ(*lacZ)M15)* | Dunn *et al.,* 2005 |
| **Plasmids** | **Relevant characteristics** | **Reference** |
| pEVS104 | conjugative helper, *oriV_R6k_*_γ_, *oriT, Kn^R^* | Stabb and Ruby 2002 |
| pVSV105 | *oriV_R6k_*_γ_, *oriV_pES213_, oriT, Cm^R^* | Dunn et al., 2006 |
| pNL6 | ES114 *hns* complement plasmid made with pVSV105*,* *oriV_R6k_*_γ_, *oriV_pES213_, oriT, Cm^R^* | Lyell et al., 2010 |
| pEVS170 | mini-Tn5 delivery vector maintained in Rho3, *oriV_R6k_*_γ_, oriT, Erm^R^, | Lyell *et al.,* 2008 |
| pLostfox-Kn | *tfoX* expression vector, *oriT*, *f1 ori*, Kan^R^ | Brooks *et al.,* 2014 |

**Biofilm Assays**. Wrinkled colony and liquid pellicle formation assays were adapted from Thompson et al., 2018.

 	Wrinkled colony assay**.** Strains were streaked out on LBS or LBS + erm agar plates and grown overnight at 24°C. Single colonies were then cultured in 5 mL LBS broth shaking (200 rpm) overnight at 24°C. The next day, they were subcultured in 1:80 ratio and let grow to early log phase. The optical density at 600 nm (OD_600_) was normalized to 0.2 using LBS broth, and 10 uL of the culture were spotted on LBS agar plates three times. After incubating spots for 48 hours at 24°C, these were imaged under white light using the Leica M165 FC dissecting microscope with the Flexacam C3 camera.

 	Pellicle formation assay. Strains were grown and cultured as described above. The subcultures were grown to mid-log phase, and the optical density at OD_600_ was normalized to 0.2 using LBS broth in a volume of 2 mL. The normalized cultures were then added to a 12-well microtiter plate and incubated statically at 24°C for 48 hours. The microtiter wells were imaged under white light using the Leica M165 FC dissecting microscope with the Flexacam C3 camera. At the end, pellicles were disrupted with a pipette tip and imaged once more.

**Squid single strain colonization assay.** *E. scolopes* hatchlings were colonized with approximately 7x10^2^-10^4^ CFU/mL of bacteria according to standard procedure (Naughton and Mandel 2012). At 48 hours post-inoculation (hpi), luminescence of hatchlings was measured using the Promega GloMax 20/20 luminometer and hatchlings were euthanized by storage at -80°C. CFU counts per light organ were determined by plating homogenized euthanized hatchlings onto LBS plates and counting colonies.
